# Cognitive Flexibility and Brain Network Energy in Healthy Aging: an Allostatic Perspective from the SENECA model

**DOI:** 10.1101/2025.05.16.654421

**Authors:** Clément Guichet, Sophie Achard, Martial Mermillod, Monica Baciu

## Abstract

Understanding how the older adult brain sustains cognitive flexibility remains a central question in aging research. Here, we analyzed resting-state fMRI data from the population-based CamCAN database (N = 628; age 18–88) and applied structural balance theory to measure functional network energy, a graph-theoretical proxy of network flexibility. In line with the SENECA model, our findings highlight midlife as a critical transition period: network energy is redistributed along the sensory-transmodal hierarchy, shifting from higher-level networks (DMN–FPN) to lower-level networks (SMN, CON, Auditory, Visual, Language). This reorganization (i) helps preserve a global wiring economy across the lifespan, hinting at an allostatic mechanism (i.e., stability through change); and (ii) may support embodied semantic strategies in older adulthood, leveraging more predictive processing to sustain cognitive flexibility at lower costs. Taken together, our study reframes healthy neurocognitive aging as an allostatic process and provides a reference for extending the SENECA model to metabolism and neuropathology.

## 1. Introduction

As the global population ages (United Nations, 2023), understanding the cerebral mechanisms of healthy aging has become a pressing goal to inform interventions that may mitigate cognitive decline. Among the many facets of cognition, *cognitive flexibility* – the ability to adapt thinking and behavior to changing goals and contexts – stands out as a central determinant of optimal cognitive functioning across the lifespan (Politakis et al., 2022; Uddin, 2021), with applications in language, mathematics, perception, and rule use (Ionescu et al., 2024). Yet, how the aging brain sustains cognitive flexibility remains insufficiently understood.

This study addresses this question through the lens of language production, focusing on interactive processes such as semantic control. Below, we argue that semantic control provides a unique bridge between cognitive flexibility and cognitive aging, and propose to integrate these perspectives by testing predictions of the SENECA model (Guichet, Banjac, et al., 2024).

### Semantic control : a window into cognitive flexibility

A major challenge in studying cognitive flexibility across the lifespan is the lack of conceptual clarity regarding its definition (Ionescu, 2022; Morra et al., 2025). A central debate opposes views of flexibility as a “*domain-general executive resource*” (Bialystok & Craik, 2006; Diamond, 2013; Miyake et al., 2000) versus a “*domain-specific property*” (e.g., language flexibility, mathematical flexibility) (Ionescu, 2012, 2017). Recently, a more interactive account proposes that flexibility may represent a core mechanism “*duplicated*” across cognitive domains, making it neither strictly domain-general nor domain-specific, but the product of both (Endress, 2019; Ionescu et al., 2024).

Language offers a particularly powerful framework for examining this view. For example, the Language-union-Memory model (L∪M) (Roger et al., 2022) conceptualizes language as the interplay between domain-general control and domain-specific lexico-phonological and semantic memory systems. Similarly, Bourguignon & Lo Bue (2025) describe language as an emergent property of interactions among articulatory-rehearsal, executive, and semantic components. Within this perspective, semantic control – the ability to selectively access and manipulate learned semantic knowledge (Reilly et al., 2023) – appears as key expression of cognitive flexibility through language-memory-executive interactions.

This is made more evident at the brain network level, where semantic control and cognitive flexibility share overlapping substrates, primarily engaging the semantic-oriented *Default Mode Network* (DMN) and the control-oriented *Fronto-Parietal Network* (FPN) (Branzi & Lambon Ralph, 2023; Chiou et al., 2023; Kupis et al., 2021; Nieberlein et al., 2024; Wang et al., 2024; for a meta-analysis on flexibility, see Xia et al., 2024). More specifically, the DMN supports semantic memory and self-referential processes (Menon, 2023; Raichle, 2015), whereas the FPN underpins goal-oriented control (Camilleri et al., 2018). In sum, this overlap in brain architectures further elevates semantic control a key window into the neural mechanisms of cognitive flexibility, and more broadly, into how the brain explores and regulates semantically grounded, goal-directed behaviors (Khajehabdollahi et al., 2025).

### Semantic control : a window into cognitive aging

Semantic control also offers a compelling framework for understanding cognitive aging (Benítez-Burraco & Ivanova, 2024), as it bridges the two dominant trajectories with age: stable or enriched semantic knowledge and declining executive control (Loaiza, 2024; Park et al., 2001; Salthouse, 2019). While older adults typically show expanded semantic stores and preserved comprehension (Alves et al., 2021; Cutler et al., 2025; Diaz et al., 2016; Harada et al., 2013), they experience reduced inhibitory control, leading to greater difficulty regulating lexico-semantic access. This may manifest as more frequent tip-of-the-tongue experiences or prolonged naming latencies, particularly beyond midlife (Benítez-Burraco & Ivanova, 2023; Hoffman & MacPherson, 2022; Oosterhuis et al., 2023; Shafto & Tyler, 2014; Verhaegen & Poncelet, 2013).

At the brain level, the DECHA model (Spreng et al., 2018; Spreng & Turner, 2019) proposes that reduced inhibition primarily impairs DMN deactivation, yielding more rigid DMN–FPN coupling with age. This rigidity is thought to induce significant semantic control challenges from midlife onwards, as persistent DMN activation introduces interference during lexico-semantic access (Guichet, Banjac, et al., 2024; Krieger-Redwood et al., 2019; Martin, Saur, et al., 2022; Martin, Williams, et al., 2022).

### The SENECA model

Building on this, recent theories suggest that midlife marks a broader neurocognitive transition from exploration to exploitation (Spreng & Turner, 2021) or from learning to prediction (Brown et al., 2022), as cognition becomes increasingly grounded in accumulated semantic knowledge (i.e., persistent DMN activation) and less responsive to high-control demands (more inflexible coupling with the FPN). Yet, the mechanistic basis of this transition remains unclear.

The SENECA model (Guichet, Banjac, et al., 2024) addresses this gap, proposing that the transition in midlife is part of an allostatic process. Allostasis refers to the predictive regulation of energetic resources (Katsumi et al., 2022), providing stability of the system through change (Theriault et al., 2025). In the context of healthy aging, allostasis may refer to brain network adaptations with the aim of preserving a *global wiring economy* – supporting efficient information transfer at a minimal wiring cost across the lifespan (Achard & Bullmore, 2007; Bullmore & Sporns, 2012). Specifically, we proposed that these network adaptations involve a shift from long-range, resource-intensive, DMN-FPN dynamics to a short-range, energy-efficient, lower-level networks architecture involving the SMN and the CON (Guichet, Banjac, et al., 2024). Converging evidence supports this view, showing that the age-related vulnerability of high-level associative systems, such as the DMN and FPN, stems from their high energetic demands (Deery et al., 2024; Diniz & Crestani, 2023; Leonards et al., 2023; Stiernman et al., 2021), due in large part to their long-range connectivity (Li et al., 2023; Luppi, Sanz Perl, et al., 2024). Specifically, the posterior cingulate cortex (PCC) – a key DMN-FPN hub (Krieger-Redwood et al., 2016) with high metabolic constraints (Basten et al., 2015; Utevsky et al., 2014) – has been identified as the main driver of this vulnerability (Godbersen et al., 2023; Hahn et al., 2024). Altogether, these findings suggest that midlife likely represents a pivotal adaptation period akin to an allostatic regulation: the brain adapts its “*resource allocation strategy*” in face of gradually declining metabolic resources (Deery et al., 2024; Ramchandran et al., 2019) to maintain a global wiring economy.

Guided by the SENECA model, this study examines how brain functional energy, which we later interpret as network flexibility (see Method), relates to cognitive flexibility, which we operationalized via semantic control.

1. **Whole-brain level** – We predict stable overall network energy and global efficiency across adulthood, consistent with a preserved global wiring economy.
2. **Subnetwork level** – To maintain such wiring economy, we expect a reallocation of energy around midlife: from DMN-FPN dynamics, particularly engaging the pmDMN and supporting semantic control in youth, toward lower-level networks, emphasizing the SMN and the CON and potentially supporting enhanced semantic access in older age.
3. **Region level** – Given the heterogenous integration of the PCC within the DMN (Godbersen et al., 2023), we anticipate a reallocation of energy from the dorsal (dPCC) component – linked to FPN coordination – toward the ventral (vPCC) component, associated with self-referential processes (Leech et al., 2011; Leech & Sharp, 2014).

## 2. Material and Methods

### 2.1. Participants

Our dataset included 628 healthy adults (age range: 18-88; 310 males, 318 females) from the Cambridge Center for Ageing and Neuroscience Project (http://www.mrc-cbu.cam.ac.uk/: Cam-CAN et al., 2014). All participants gave written informed consent following approval by the Cambridgeshire 2 Research Ethics Committee (reference: 10/H0308/50). Details on participant recruitment are provided in Taylor et al. (2017).

This sample size was obtained by including the participants with at least 5 (out of 8) cognitive scores. Any missing score was imputed with the median value of the participant’s age decile. These 8 scores were derived from neuropsychological tasks assessing semantic control performances both directly and indirectly (see Table 1 below).

**Table 1.** Neuropsychological tasks and associated cognitive processes were considered in this study. A detailed description can be found in Supplementary information.

| Neuropsychological test | Cognitive processes | Task type and relation to semantic control |
| --- | --- | --- |
| **Picture naming** | Language (phonological and semantic access) | Direct |
| **Verbal fluency** | Language (phonological and semantic access) and EF | Direct |
| **Tip-of-the-tongue situations** | Language (retrieval) and EF (monitoring) | Direct |
| **Proverb comprehension** | Language (comprehension) and EF (abstraction) | Language, Indirect |
| **Sentence comprehension** | Language (syntax, semantic) | Language, Indirect |
| **Story recall** | Long-term memory | Language, Indirect |
| **Cattell task** | EF (Logical reasoning, problem-solving) | Domain-general, Indirect |
| **Hotel task** | EF (multitasking, planification, self-monitoring) | Domain-general, Indirect |

### 2.2. MR acquisition and preprocessing

*MR acquisition*. Resting-state functional MR images were acquired on a 3T Siemens TIM Trio with a 32-channel head coil for all participants at the Medical Research Council and Brain Sciences Unit in Cambridge, UK (MRC-CBSU). Two hundred and sixty-one volumes (261) were acquired with eyes closed in descending order using a gradient echo-planar imaging sequence lasting 8 min and 40 s (GEEPI, 32 axial slices, 3.7 mm thickness and interslice gap of 20% for whole brain coverage including cerebellum, TR = 1.97 ms, TE = 30 ms; voxel-size 3 x 3 x 4.44 mm, flip angle = 78°, field of view = 192 x 192 mm). In addition, structural images were acquired using a 1mm3 isotropic, T1-weighted Magnetization Prepared RApid Gradient Echo (MPRAGE) sequence and a 1mm3 isotropic (more information about the MR acquisition and resting state protocol are provided by Cam-CAN et al., 2014).

Data were preprocessed using the *fMRIPrep* software (https://fmriprep.org/en/stable/: Esteban et al., 2019). T1-weighted images underwent skull stripping, tissue segmentation, and spatial normalization. Resting-state fMRI data underwent motion correction, slice timing correction, susceptibility distortion correction, co-registration, and spatial normalization.

#### Functional connectomes

Denoised timeseries were aggregated using the 360-region HCP-MMP 1.0 atlas (Glasser et al., 2016).We used custom scripts to co-register the atlas to subject-space with ANTs (https://github.com/ANTsX/ANTsPy) and applied a confound removal strategy via Nilearn (https://nilearn.github.io/). Confounds included high-pass filtering, the full 24 motion parameters, and the full 8 white matter and CSF parameters as recommended in Wang et al. (2024). No global signal regression was performed as it introduces artefactual negative connections (Chai et al., 2012; Murphy & Fox, 2017; Saberi et al., 2022). We performed full scrubbing with a framewise displacement threshold set to 0.5 mm and the standardized DVARs to 1.5. In the event that all segments were removed with full scrubbing, the scrubbing parameter was decremented. Mean framewise displacement was also extracted and used as a covariate in subsequent statistical models. Spatial smoothing and standardization was applied to the signal using a 6mm full-width-at-half-maximum (FWHM) kernel. Finally, functional connectomes were obtained by computing the Pearson correlation between time series.

### 2.3. Network Energy

#### 2.3.1. Network energy function

To operationalize the concept of brain network flexibility, we leveraged *structural balance theory* and the concept of network energy – a graph-theoretical proxy for brain functional dynamics initially developed by Marvel et al. (2009) and recently applied to brain networks by Saberi et al. (2024). This framework provides a principled way to examine the role of signed network motifs, which are often disregarded in conventional graph-theoretical approaches (e.g., negative weights set to zero). By quantifying the relative prevalence of balanced versus imbalanced motifs, the energy function offers a tractable approximation of flexibility in functional brain networks.

The network energy of signed and fully-weighted and signed graph in Equation 1. All computations were performed in Python (version 3.9.12).

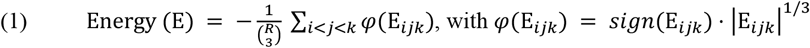

With E_*ijk*_ = *W*_*ij*_*W*_*ik*_*W*_*jk*_ representing the product of the pairwise Pearson correlations (*W*) in the motif defined by regions *i, j*, and *k*. Each *W* can be positive or negative, thereby representing correlated or anti-correlated function activity. The product of each motif defined by *φ*(E_*ijk*_) can be either positive, indicating a balanced motif with an even number of anti-correlations, or negative, indicating an imbalanced motif with an odd number of anti-correlations. The final network energy metric (E) is obtained by taking the average of all motifs in the network and applying the negative sign in front of the equation.

In sum, positive values of network energy indicate a majority of imbalanced motifs in brain network organization, which can be interpreted as flexibility. Conversely, negative values indicate a majority of balanced motifs, which can be interpreted as rigidity (Figure 1).

**Figure 1.**
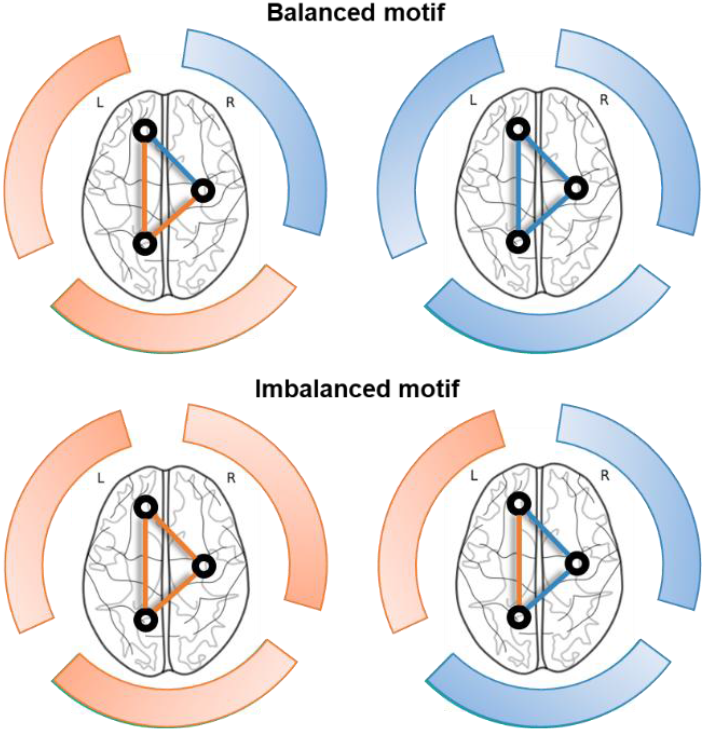
Network energy framework. Based on the framework described in Saberi et al. (2024). Blue and Orange represent correlated and anti-correlated patterns of functional connectivity respectively

#### 2.3.2. Subnetwork level

We further examined the energy *within* networks and *between* networks. Within networks, we examined all motifs made up of regions from a single network N (see Equation 2a).

*Let N*_*α*_ *represent a single network, and let I*_*α*_ *be the set of indices for regions in network N*_*α*_.

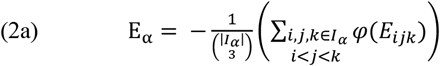

Between networks, we examined motifs made up of at least one region from each network N (see Equation 2b). This allowed us to examine the motifs that resume the joint connectivity between pairs of networks.

*Let N*_*α*_ *and N*_*β*_ *represent two networks, and let I*_*α*_ *and I*_*β*_ *be the sets of indices for regions in networks N*_*α*_ *and N*_*β*_ *respectively*.

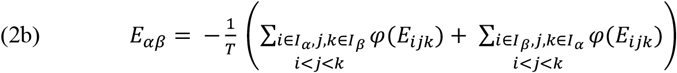

With 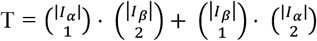 representing the two ways to select motifs spanning both networks: (i) a motif made of one region from N_*α*_ and two regions from N_*β*_; and (b) a motif made of one region from N_*β*_ and two regions from N_*α*_.

*Between-Within network*. We quantified the tendency of a network to exhibit more energy between versus within networks. We begin by making all values positive to avoid inconsistencies (Equation 3a). Then, we compute the average energy of all between and within components (Equation 3b & 3c) and their normalized ratio (Equation 3d).

*Let* E_*within*_ *be the set of all within-network energy values*, E_*between*_ *that of all between-network values, and* E_*all*_ *their union*.

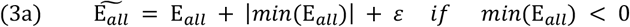

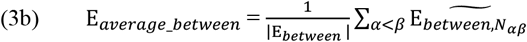

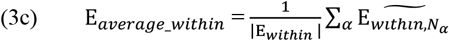

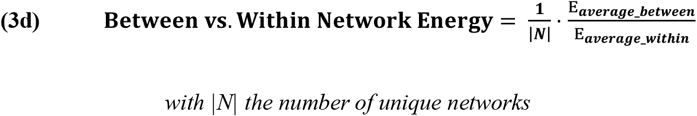

*with* |*N*| *the number of unique networks*

#### 2.3.3. Region level

Based on the Local Balance Index (Diaz-Diaz et al., 2024; Khanafiah & Situngkir, 2004), we computed the ***Region Energy (RE)***. It quantifies the ratio of balanced versus imbalanced motifs to which a given region contributes, indicating whether a region drives flexibility (RE > 1) or rigidity (RE < 1) in the network.

We first define the signed contribution of a region *r* as follows:

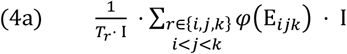

With 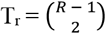 is a normalizing factor, representing the number of motifs involving region *r*. I is an indicator function observing whether the motif (E_*ijk*_) is strictly negative or positive. Then, we defined the ***Region Energy*** as:

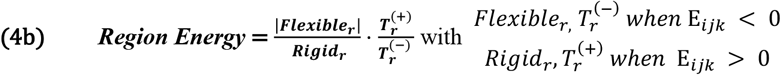

### 2.4. Statistical analysis

Statistical analyses were carried out in R (version 4.3.2). All models included sex, handedness, total intracranial volume, and mean framewise displacement as covariates.

#### Whole-brain level

We computed the whole-brain network energy (see Equation 1) for each individual and modeled its relationship with age using a Bayesian linear regression model implemented in the *brms* package (version 2.22.0; Bürkner, 2017). The model was fitted using four chains, each with 4,000 iterations and 2,000 warmup steps, with hyperparameters adjusted to ensure proper convergence. Using the *bayestestR* package (version 0.13.2; Makowski et al., 2019), we conducted a practical equivalence test (Kruschke, 2018) with a 89% highest density interval (HDI) to test whether whole-brain energy remained stable across the lifespan. The region of practical equivalence (ROPE) limits were defined automatically using the *rope_range()* function. We also computed the average between-vs. average within-network energy (see Equation 3) and fitted a standard frequentist GAM with age as a smooth predictor (parameters: *k = 3, method = REML*) to investigate subnetwork energy tendencies across the lifespan.

#### Subnetwork level

We computed the energy within each network and between pairs of networks (see Equation 2) for 9 of the 12 networks mapped by Ji et al. (2019). We did not consider the lesser known networks (i.e., ventro-multimodal, posterior-multimodal, and orbito-affective). Then, we followed the same GAM procedure to examine the age effect on each network or pair of network. Significance thresholds were adjusted at the False Discovery Rate (FDR) across models that examined similar neural elements (that is either within-network or between-network).

#### Region level

We computed the energy for each DMN and FPN regions with respect to DMN-FPN motifs (see Equation 4), thus indicating how each region drives flexible or rigid DMN-FPN dynamics. Similarly, we computed the energy of posterior-medial (pm) DMN regions only considering DMN motifs. pmDMN regions were taken from the Posterior Cingulate division of the HCP-MMP1.0 atlas (Glasser et al., 2016). Then, we followed the same GAM procedure to examine the age effect as described previously.

### 2.5. Null models

We sought to replicate our analyses using thresholded matrices to minimize the number of functional connections caused by noise. We determined the statistical significance of each correlation coefficient following two approaches. In the first approach, we generated surrogates by randomizing timeseries, thereby preserving spatial and temporal autocorrelation as these measures are very sensitive to age in our dataset (Shinn et al., 2023). In the second approach, we determined an absolute threshold (± 0.27) through the correlation screening method, with the goal of limiting false-positives due to intra-regional correlation (Lbath et al., 2024).

We added the subject-specific density as a covariate to preclude the effect of different network sizes between individuals (Rubinov & Sporns, 2010; Stumme et al., 2022; van den Heuvel et al., 2017). Indeed, the elimination of non-significant edges may distort graph theoretical measures due to differences in the number of remaining edges, and generally lead to more random network characterization in network in older adults due to overall low FC.

### 2.6. Neurocognitive analysis

To examine the relationship between region energy and semantic control across the lifespan, we employed Partial Least Squares (PLS) correlation analysis using the toolbox myPLS (https://github.com/MIPLabCH/myPLS) in MATLAB R2020b. PLS is a statistical technique designed to identify latent relationships between brain features (*X matrix*), here the region energy values obtained via the spatiotemporal null models (section 2.5), and cognitive (*Y matrix*) performances.

#### Data preparation

For the cognitive matrix, we included the 8 cognitive scores presented in Table 1 as well as two variables derived from piecewise polynomials. These polynomial functions allow the model to optimize the covariance between energy values with potential nonlinear age trajectories. The first function represents an acceleration after a transition age *t*, and complementarily the second represents a leveling-off after *t*. We fixed the transition age at age 53 in line with results reported in section 3. This approach has been validated in previous teamwork (Guichet, Harquel, et al., 2024; Guichet et al., 2025).

#### PLS inference

Briefly, we computed the covariance matrix (X*Y^T^) and performed singular value decomposition (SVD) to extract latent components. Each component includes singular values (S) reflecting shared information, and brain (U) and cognitive (V) saliences reflecting feature contributions. Statistical significance and robustness were assessed via 10,000 permutations and 1,000 bootstrap resamples, respectively. Features with a bootstrap ratio (BSR ± 3) – the salience weight divided by its bootstrapped standard deviation – were considered robust at the 99% confidence level (Krishnan et al., 2011).

## 3. Results

### 3.1. Whole-brain level

In line with our hypothesis, a Bayesian practical equivalence test indicated suggests no change in whole-brain network energy at rest throughout the adult lifespan (see Figure 2A) : the ROPE (Region of Practical Equivalence) fully covered the 89% HDI of posterior samples [-1e-4; 1e-4]. Among covariates, only gender showed a significant effect, with women exhibiting higher network energy than men [0.01; 0.02]. In parallel, energy was increasingly reallocated towards within-rather than between-network connectivity with age, with an inflection point at age 52 (*F* = 78.01, *p* < .001, *edf* = 1.93, signed partial R^2^ = -0.2) (see Figure 2C).

**Figure 2.**
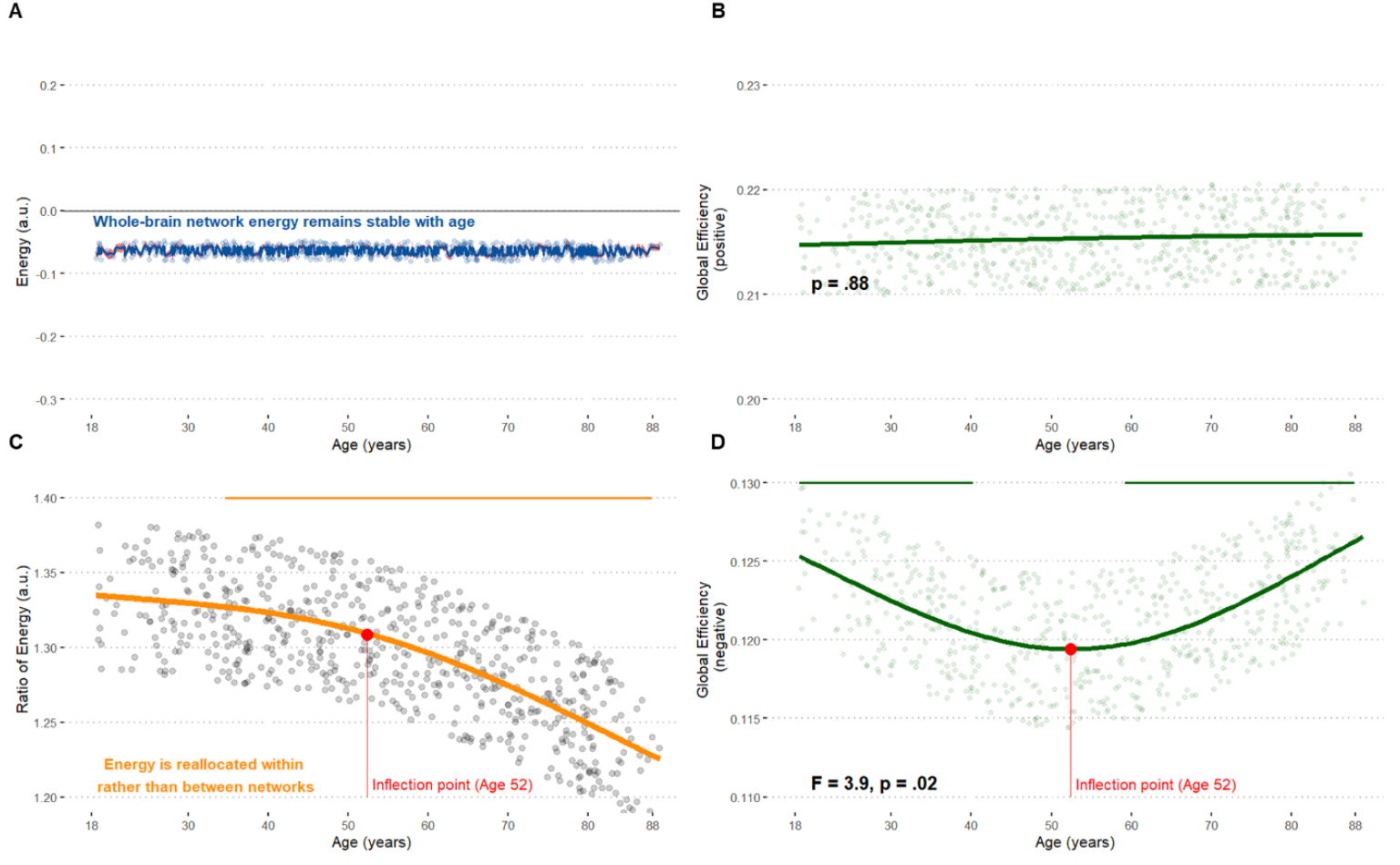
Whole-brain analysis. **(A)** Mean of the posterior predictive distribution from the Bayesian model. **(B & C & D)** Age-related trajectories from GAMs. The horizontal bars on top indicates the age window of significant changes, determined from the 1^st^ derivative of the GAM smooth function. The red point marks the inflection point, corresponding to the age with the largest significant 2^nd^ derivative

Complementary analyses indicated that this reallocation was primarily driven by anticorrelated dynamics: Positive (correlated FC) global efficiency remained stable across the lifespan (*p* = .88, Figure 2B), whereas negative (anticorrelated FC) global efficiency showed a similar inflection at age 52 (*p* = .02; Figure 2D).

### 3.2. Subnetwork level

We decomposed whole-brain energy (i) within networks, and (ii) between networks (Figure 3). Model statistics are reported in Table S1 in Supplementary information. Overall, our results clarify the previous tendency, indicating that the reallocation of energy around midlife benefits lower-level networks, with a similar inflection point at age 52.

**Figure 3.**
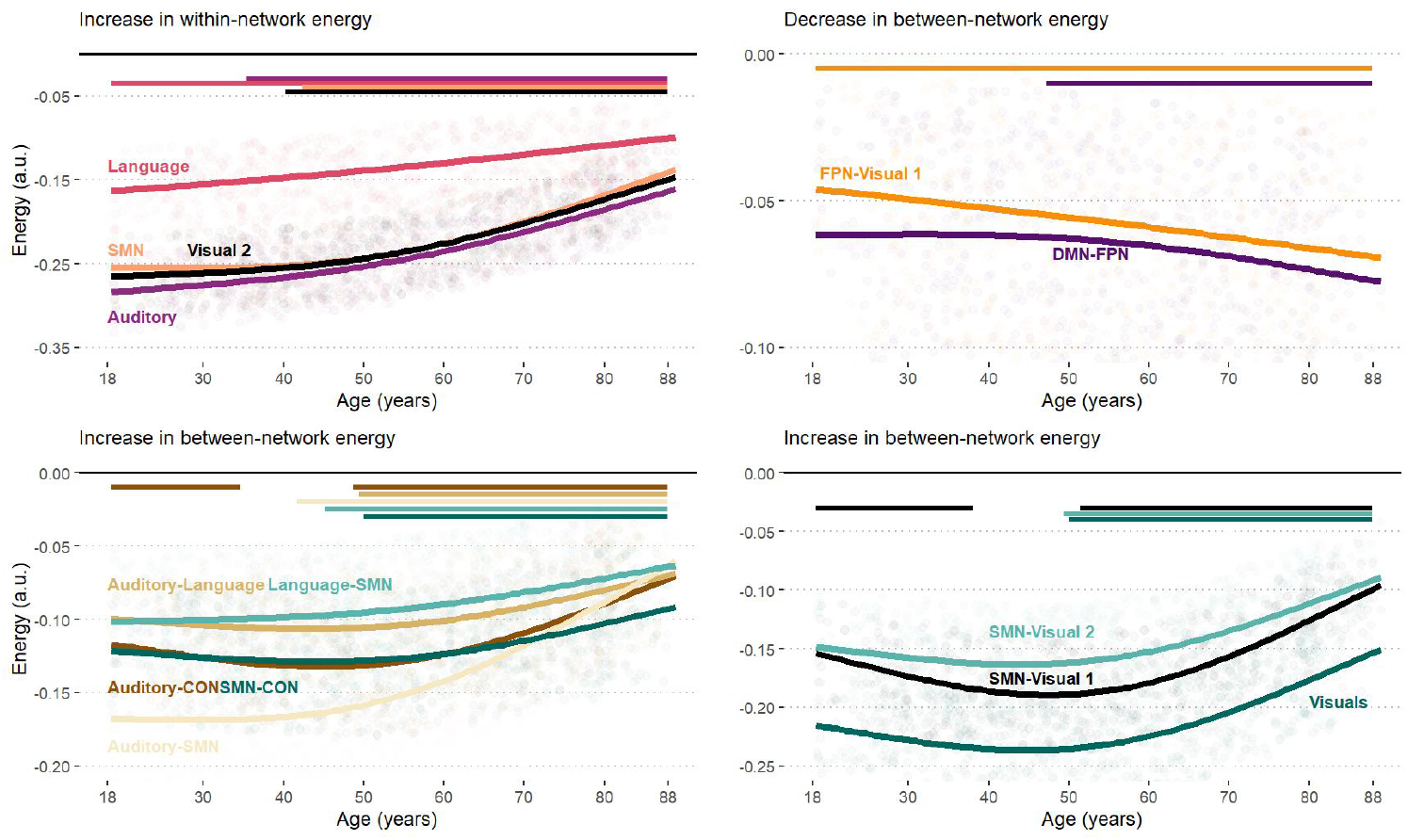
Age-related within and between-network energy. The trajectories plots the predicted values of Generalized Additive Models (GAM). Bars on top represent the range of significant age-related changes and were determined by computing the 1^st^ derivative of the fitted GAM smooth function. Only the largest trajectories are displayed

#### Within networks

Age-related changes occurred within lower-level networks, especially within the SMN (partial R^2^ = .11), Auditory (.10), Language (.09), and Visual 2 (.09) networks (see upper left panel on Figure 3). *Between networks*. Only DMN-FPN had a substantial decline in network energy beyond midlife (partial R^2^ = -.04) (see upper right panel on Figure 3); whereas energy across lower-level networks exhibited important increases, especially considering Auditory-SMN (.14), Auditory-CON (.09) and SMN-Visual couplings (.05) (see bottom panels on Figure 3).

### 3.3. Region energy in DMN-FPN & within DMN

#### DMN-FPN

In line with our hypothesis, Figure 4A shows that the decline in DMN-FPN network energy is primarily driven by posterior cingulate regions, especially bilateral areas 7m and the bilateral parieto-occipital sulci (POS2). Overall, regions with stronger contributions were more posterior (*r*_*(76)*_ *=* .52 [.34;.67] 95% CI, *p* < .001), superior (*r*_*(76)*_ *=* .32 [.11;.51] 95% CI, *p* < .001), and medial (*r*_*(76)*_ *=* .32 [.1;.5] 95% CI, *p* < .001).

**Figure 4.**
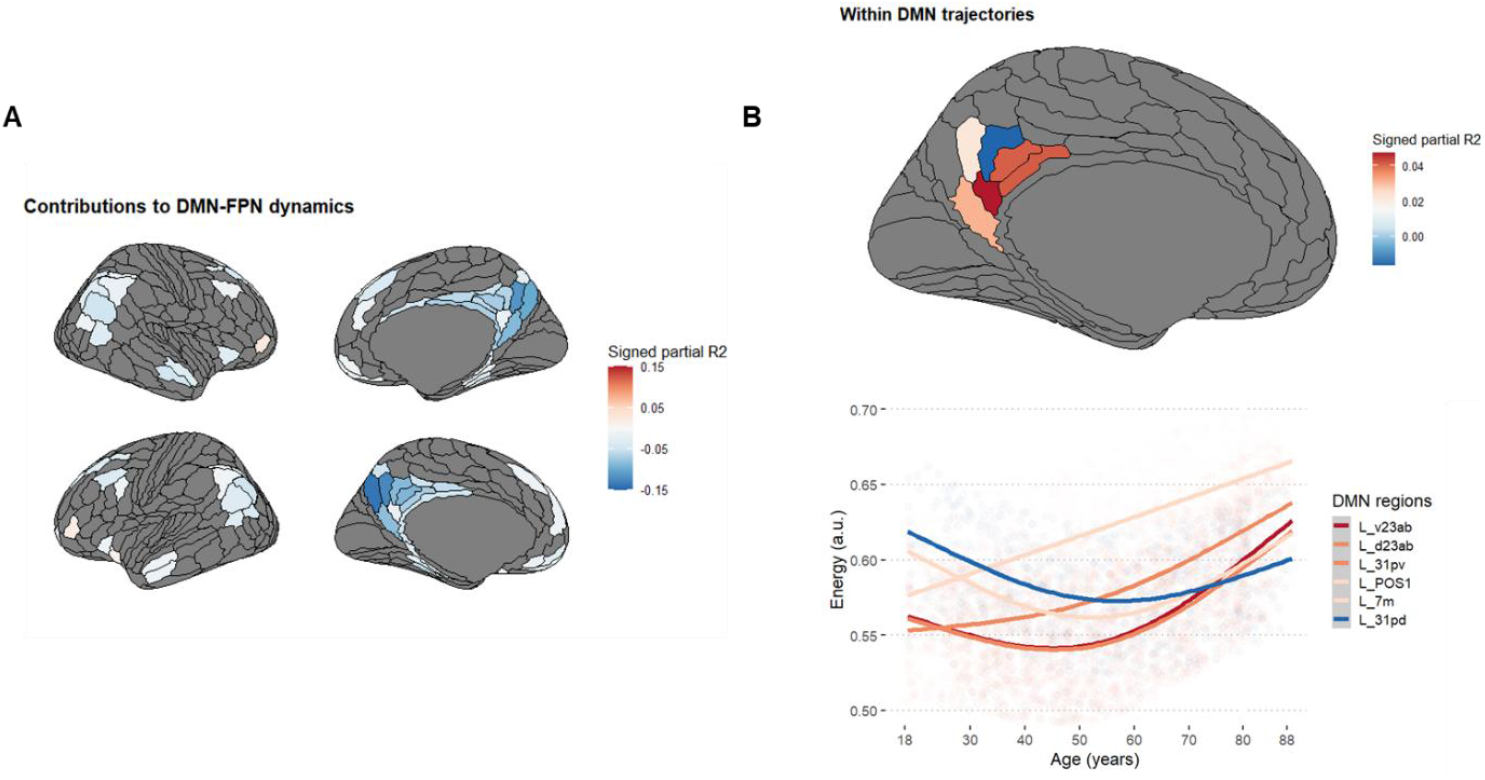
Regions contributing to changes in (A) DMN-FPN and (B) within DMN energy.

#### Within the DMN

Figure 4B shows heterogeneity of the PCC activity within the DMN, but only in the left hemisphere. The left dPCC showed substantial decline in energy until midlife, indicating more rigidity, at which point the left vPCC showed increased energy, indicating more flexibility.

### 3.4. Region energy at the whole-brain level

Next, we evaluated the contribution of each region to imbalanced versus balanced motifs. In line with our previous results, Figure 5 illustrates that most changes have an inflection point in midlife, with more energy circulating in sensorimotor, auditory and visual cortices; and less so in mostly posterior DMN and FPN areas. A thousand spin tests preserving spatial autocorrelation revealed that these changes were strongly and significantly correlated to changes along the sensory-transmodal (G1) (*r*_*s*_ = .49, *p*_*spin*_ < .001).

**Figure 5.**
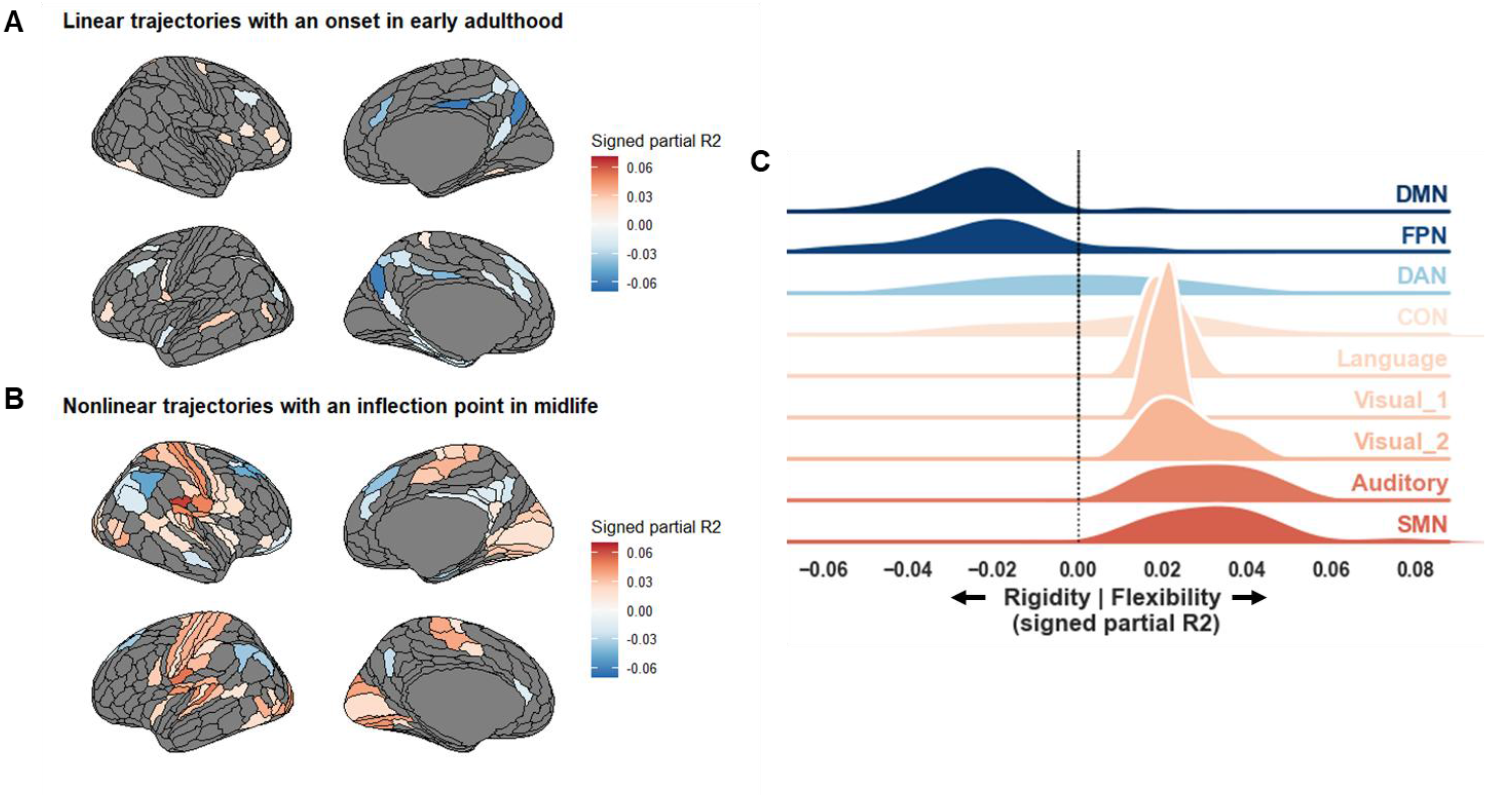
Age-related changes in Region Energy. **(A & B)** Brain illustrations. Red indicates increased energy, pointing towards more flexibility, and Blue indicates decreased energy, pointing towards rigidity. **(C)** Resting-state network description of panel A & B

These findings remain valid when thresholding connectivity matrices using null models that preserve spatial and temporal autocorrelation (Shinn et al., 2023). When applying the more stringent correlation screening approach (Lbath et al., 2024), whole-brain and network-level results result also held. However, robustness could not be assessed at the region level using this approach as there was not enough significant connections to conduct reliable analyses (see Supplementary information).

### 3.5. Neurocognitive analysis

We further sought to relate region energy changes to cognitive flexibility, through the lens of semantic control performances. Partial Least Squares (PLS) correlation analysis revealed one major cognitive control component (LC1) which explained 80.7% (*p*_*FDR*_ = .001) of the total shared variance and one smaller semantic component (LC2) which explained 8.9% more variance (*p*_*FDR*_ = .031). Below, we detail the brain-cognition relationships for each.

#### 3.5.1. Cognitive control – Latent component 1

At a cognitive level, LC1 is strongly associated with fluid intelligence (Cattell task ; 14.59), picture naming (13.55), multitasking (7.44), reduced tip-the-tongue occurrences (7.36), and enhanced verbal fluency (7.12). Performance along this component is associated with high energy in DMN and FPN regions alongside low energy sensorimotor and visual cortices (see Table S2, Supplementary information). This brain-cognition relationship follows a nonlinear age-related decline, with a notable reversal around age 60 (i.e., crosses zero) which could reflect the reallocation from higher to lower-level networks reported in section 3.2 and 3.4.

#### 3.5.2. Semantic – Latent component 2

LC2 is strongly associated with verbal fluency (9.72), long-term memory (7.37), proverb comprehension (4.73), and picture naming (3.33), with a negative association with tip-of-the-tongue occurrences (-3.87), indicating that this component reflects semantic processing rather than lexical retrieval and monitoring (see Table 1, section 2.1 for a description of processes of each task).

Age trajectories for LC2 show that these cognitive abilities peak in midlife and remain relatively stable into older age. However, the relationship between cognition and brain network energy is weaker. Neural patterns show a linear reduction of energy in fronto-cingulate-sensorimotor systems, including the bilateral auditory cortices, SMN and CON networks (see Table S2, Supplementary information). Note that the confidence interval hovers around zero, likely reflecting high inter-individual variability that might be captured by other age-invariant factors. In summary, LC2 reflects acute semantic access that is relatively preserved with age, supported by a modest and highly variable loss of network flexibility in a distributed lower-level brain system.

## 4. Discussion

In this study, we examined how the older adult brain reorganizes its functional architecture and the consequences on semantic control, an interactive language process between cognitive control and semantic knowledge thought to reflect cognitive flexibility. To this end, we tested the predictions of the SENECA model (Guichet, Banjac, et al., 2024) using cross-sectional resting-state fMRI data from 628 healthy adults aged 18-88. We focused on functional network energy levels, a graph-theoretical measure proposed as a proxy for brain network flexibility (Saberi et al., 2024).

Our results are consistent with our hypotheses : (1) At the brain level, global network energy levels remains stable across the lifespan, yet midlife represents a pivotal transition period. This transition involves substantial energy reallocation in midlife from higher-to lower-level networks, likely reflecting shifts in anticorrelated dynamics. (2) At the cognitive level, this midlife reconfiguration correlates with the emergence of semantic control challenges. In older adulthood, we further observed a low-energy fronto-cingulo-sensorimotor system, potentially tied to an embodied semantic strategy for sustaining cognitive flexibility through alternative neural mechanisms.

Taken together, our study supports a view of healthy cognitive aging as an outcome of allostatic regulation: around midlife, functional brain resources are reallocated to maintain a global wiring economy within the brain, compromising on higher-level cognitive functions such as those underpinning semantic control.

### Conceptualizing healthy cognitive aging as an allostatic process

Whole-brain energy remains remarkably stable across the adult lifespan, reflecting a balance between rigidity and flexibility (see Figure 2). Saberi et al. (2024) previously interpreted this low-energy resting state as the brain’s attempt to “*minimize energy […] while maintaining flexibility*”. This view aligns with our hypothesis of a preserved wiring economy with age, suggesting an allostatic process – stability through change.

In pursuing such allostasis, we observed distinct network configurations in younger and older adulthood, consistent with prior evidence that energy minimization at rest can be achieved through multiple network states (Watanabe et al., 2014). Beim Graben et al. (2019) further proposed that transitions between these low-energy states may reflect a “*homeostatic switching process*”, a notion that resonates with our interpretation of allostatic regulation.

Our findings extend this framework by showing that negative functional connectivity (FC) is central mechanism for allostatic regulation. This observation supports prior evidence implicating negative FC in the regulation of brain network energy/flexibility (Saberi et al., 2022). Across the lifespan, Saberi et al. (2021) reported a U-shaped trajectory with minimal energy/flexibility demands in early adulthood – a pattern interpreted as reduced neuroplastic needs once cognitive efficiency reaches its peak. Similarly, we identified an inflection point in global efficiency derived from negative FC around midlife (Figure 2D), and proposed a similar interpretation below.

Following Hassett et al. (2024), we propose that negative global efficiency indexes not only neuroplasticity demands but also the maturation of a cost-efficient network topology. Elevated values may signal the development of a “*central executive switching component*” underlying inhibitory control (Fox et al., 2005; Teasdale et al., 1995). Accordingly, the midlife dip in negative efficiency observed in our study may represent a transient phase of network reorganization – a “*random conformation*” (Hassett et al., 2024) during which anticorrelated dynamics facilitate allostasis, that is the maturation of alternative wiring economy mechanisms given older adulthood’s metabolic constraints (Deery et al., 2024). In summary, anticorrelated dynamics may constitute the functional scaffold of allostasis across the lifespan. By reallocating energy across subnetworks, these dynamics may adjust wiring economy mechanisms while preserving globally efficient information transfer within positively correlated networks (Figure 2A & 2B).

### Midlife : a key period for allostatic regulation

As detailed in sections 3.2 and 3.4 (Figure 3 & 5), midlife emerges as a pivotal phase of network energy reallocation, characterized by local shifts from higher-to lower-level systems. This pattern is consistent with the idea that allostatic regulation involves local, rather global, adjustments in resting-state network energy/flexibility (Beim Graben et al., 2019), as discussed above.

On one hand, the age-related reduction in energy between higher-level networks, notably between the DMN and FPN, may indicate less coordinated communication with age – an observation supporting the DECHA model (Spreng & Turner, 2019). In line with our hypothesis, complementary results from section 3.3 further highlight the posterior cingulate cortex (PCC) as central hub mediating these age-related DMN-FPN dynamics, potentially due to its unique cortical morphology of the PCC (Willbrand et al., 2022).

On the other hand, the increase of energy within/between lower-level networks suggests more flexibility in sensory systems. A potential interpretation is that older adults rely on a more redundant functional architecture, where similar information is processed across multiple sensory modalities (e.g., auditory, visual). Such redundancy could optimize for “*robustness over precision in the neural code*” (Johnston et al., 2020; Johnston & Freedman, 2023), providing robust multi-sensory representations.. Supporting this interpretation, Stanford et al. (2024) showed that as network controllability in DMN and FPN hubs declines, redundancy serves as a “*mechanism of reserve*”, promoting “*robust control of brain network and in cognitive function in healthy aging*”.

Moreover, the elevated energy levels between the cingulo-opercular network (CON) and sensory systems (SMN, Auditory) (Figure 3) underscores the CON’s central role in executive control in older adulthood (Cao & Cannon, 2021; Crittenden et al., 2016; Hausman et al., 2022), in line with Saberi et al. (2024). In line with the SENECA model, the CON could facilitate a more economical mode of attentional regulation with age, deploying control resources transiently and efficiently compared to the FPN (Cocuzza et al., 2020). More flexible CON-sensory coupling energy could also support somatosensory feedback loops in older adulthood, crucial for action monitoring and action outcome evaluation through its action-oriented subnetwork (D’Andrea et al., 2023; Dosenbach et al., 2024). In sum, the CON may establish a more embodied form of goal-directed cognition, a possibility further supported by its tight connectivity to the somato-cognitive action network (SCAN) (Gordon et al., 2023). We develop this idea of embodied cognition in a later subsection.

In summary, allostatic regulation in aging appears to rely on increased flexibility between bottom-up attentional processes (i.e., CON) and sensory networks. Enhanced redundancy (Gatica et al., 2022) and short-range connectivity (Guichet, Banjac, et al., 2024) may thus serve as functional biomarkers of allostasis, reflecting the preservation of wiring economy through adaptive network reorganization.

### Cerebral underpinnings of cognitive flexibility in aging

As shown in section 3.5, network energy reallocation directly correlates with cognitive flexibility, as indexed by semantic control performance, and unfolds along the sensory-transmodal gradient (Margulies et al., 2016). In younger adulthood, energy primarily flows between the DMN and FPN, while lower-level networks (SMN, auditory, visual, language) remain relatively disengaged/inflexible. With aging, this pattern reverses: energy becoming increasingly concentrated within sensory systems, granted them more flexibility (Figure 5).

We propose that these changes may reflect age-related structural constraints on functional dynamics. Indeed, structure-function coupling – the correspondence between anatomical connectivity and functional co-activation – is also organized along the sensory-transmodal hierarchy (Collins et al., 2024; Fotiadis et al., 2024), and has been linked to cognitive flexibility, functional diversity (Wu et al., 2020; Yeo et al., 2015), sustained attentional performance, verbal learning and retrieval (Griffa et al., 2022). Accordingly, a possible interpretation is that lower network energy/inflexibility in sensory regions signals tight coupling to underlying structural connectivity, supporting stable sensory representations “phase-locked” to the environment (Preti & Van De Ville, 2019). These reliable sensory signals could, in turn, sustain energy-demanding/flexible transmodal processes which are largely decoupled from brain structure and associated with cognitive flexibility (Rossi-Pool et al., 2021; Schwartz, 2016).

This interpretation is further supported by the similar lifespan trajectories observed between our results and prior teamwork on structural connectivity: both exhibit a nonlinear decline with an inflection point in midlife, and both correlate with the same semantic control difficulties (Figure 6). These convergences suggest that the relationship between transmodal dynamics and cognitive flexibility can be observed across multiple imaging modalities. Accordingly, age-related decline in DMN–FPN energy, i.e., more rigid DMN-FPN coupling, likely mirrors reductions in microstructural integrity (Guichet, Roger, et al., 2024), and reduced structure-function decoupling (Guichet et al., 2025).

**Figure 6.**
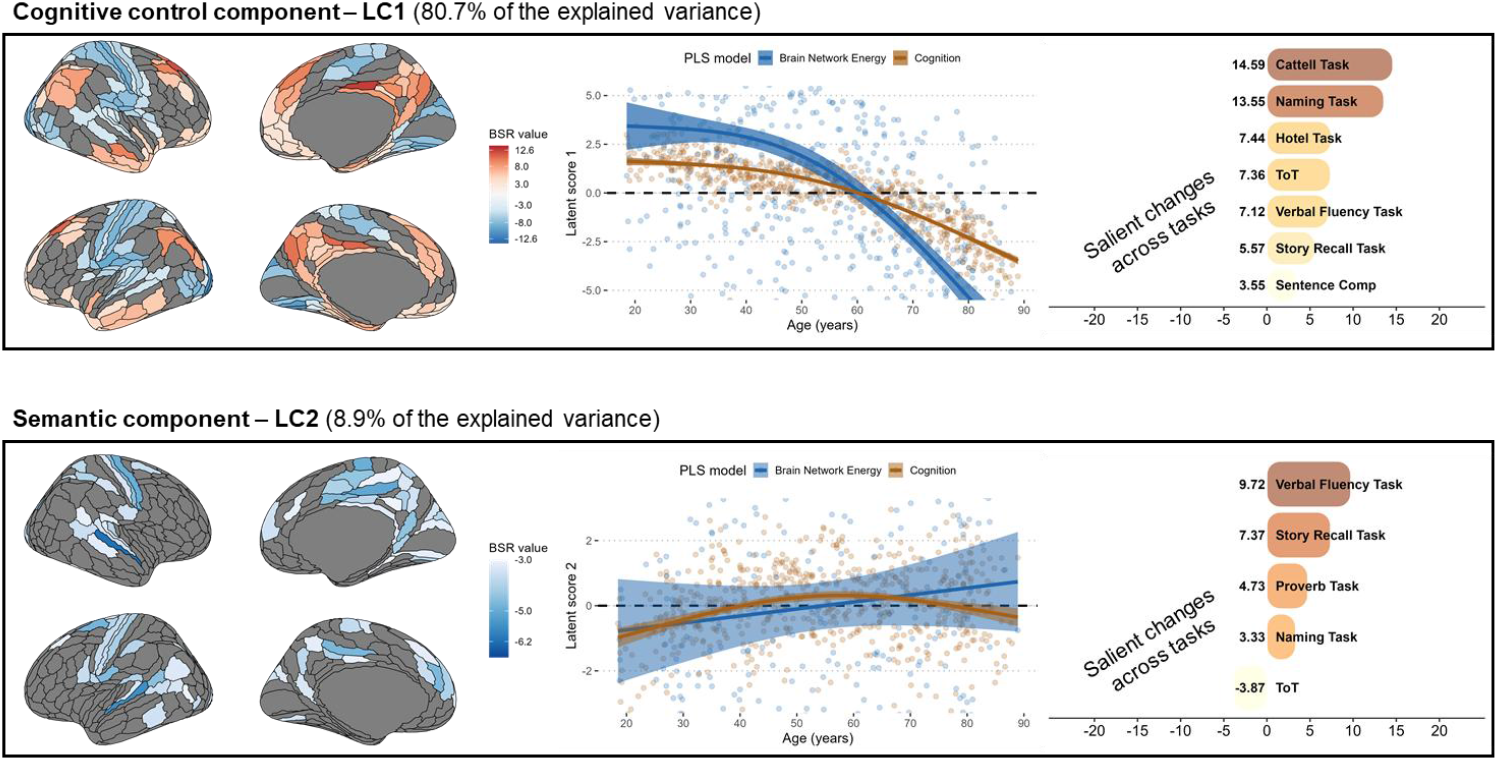
Neurocognitive analysis. The middle panels illustrate the trajectory of the first and second latent components (LC1 and LC2) inferred by the PLS. The left and right panels respectively report the brain and cognitive profiles correlated with each trajectory. Only bootstrap sampling ratios (BSR) ± 3 denote a robust contribution to the covariance patterns

### An embodied semantic strategy in older adulthood

After accounting for energy reallocation along the sensory-transmodal axis, we identified a complementary trajectory characterized by enhanced semantic access from midlife onward. This finding suggests the emergence of a *semantic strategy* in older adulthood that supports cognitive flexibility independently of executive control, consistent with its supposed protective role in language production (Baciu et al., 2021; Gollan & Goldrick, 2019; Krethlow et al., 2024).

At the brain level, this semantic strategy coincided with reduced energy/flexibility across a broad fronto–cingulo–somatosensory (FCS) system (Figure 6). This brain-cognition relationship largely overlaps with our previous findings in brain structure, which had revealed greater subcortico-sensory microstructural integrity (Guichet, Roger, et al., 2024), and stronger structure-function coupling in the SMN (Guichet et al., 2025). Together, this reinforces our interpretation that links brain energy to brain structure: reduced energy in the FCS system likely reflects a tighter alignment to the SMN’s anatomical scaffolding, effectively grounding semantic processes in sensory-driven representations.

Taken together, our study suggests that cognition becomes not only be more *semanticized* with age (Baciu et al., 2021; Spreng et al., 2018), but also more *embodied*, in line with contemporary models of semantic processing (Boleda, 2020; Borghi et al., 2024; Calzavarini, 2024; Dove, 2023; Galetzka, 2017; Hagoort & Özyürek, 2024; Kewenig et al., 2024). This embodied mechanism may operate by comparing sensory information to long-term *situation models* – semantic memory representations accumulated through a lifetime of learned associations between objects and contexts (Baltes & Lindenberger, 1997; Kintsch & Van Dijk, 1978; Lindenberger & Baltes, 1994; Lindenberger & Mayr, 2014; Martin, 2016; Wright, 2016). More broadly, this aligns with emerging frameworks that conceptualize aging as shift toward more predictive cognition (Brown et al., 2022), enabling “*the binding of sensory information towards the more general notion of semantically related episodes*” (Recanatesi et al., 2021). This is also consistent with the results reported in section 3.4 (Figure 4). In midlife, the reallocation of energy from the dPCC to the vPCC could mark a transition to a predictive mode, particularly given that the vPCC is thought to play a key role in generating predictions and the instantiation of situation models (Geerligs et al., 2022).

From an energy perspective, a predictive mode may reflect allostatic compromises towards more economical processing (Katsumi et al., 2022). Indeed, prediction appears as a resource-optimization mechanism that compresses information and minimizes energy expenditure (Elias, 1955; Price & Gavornik, 2022), others noting that it saves on metabolic costs by reduced encoding of incoming sensory signals (Katsumi et al., 2022; Theriault et al., 2021).

### SENECA: Towards an allostatic model of healthy cognitive aging

Our findings reframes healthy cognitive aging not as decline or compensation, but as an allostatic process – a reallocation of functional resources aimed at preserving a global wiring economy. The SENECA model stands in line with the “*allostatic-first*” perspective recently advanced by Theriault et al. (2025) which views cognitive decline, not as indication of malfunction, but rather as a compromise or a cost of adaptation.

Our study highlights midlife as key period for allostatic regulation. The brain appears to recalibrate its wiring economy mechanisms by modulating *anticorrelated dynamics* among three major systems: the control-related frontoparietal network (FPN), the self-referential default mode network (DMN), and the action-oriented sensorimotor and cingulo-opercular networks (SMN/CON). This transition gives rise to two distinct modes of cognitive flexibility.

In younger adulthood, cognitive flexibility primarily depends on dynamic interactions between the DMN and FPN, with the dorsal posterior cingulate cortex (dPCC) acting as a pivotal hub that coordinates domain-general executive control with language-specific semantic systems. Cooperative DMN–FPN coupling may facilitate the integration of novel experiences and goal-directed behavior, while competitive interactions help suppress irrelevant associations when top-down control is required to manage semantic complexity (Krieger-Redwood et al., 2019). This organization aligns with an *explorative or learning mode* of cognition (Brown et al., 2022; Spreng & Turner, 2021).

In the pursuit of allostasis, older adults down-regulate this DMN-FPN architecture in favor of a more sensory-driven and redundant network topology likely engaging a broad fronto-cingulo-sensorimotor configuration. This topology provides the basis for an embodied semantic strategy for cognitive flexibility, whereby lifelong semantic knowledge is efficiently compared against stable multisensory inputs, mediated by the ventral posterior cingulate cortex (vPCC). This organization aligns with greater predictive processing with reduced reliance on executive control (Bar, 2007; Bubic, 2010; Chan et al., 2021; Yang et al., 2024), consistent with previous reports of an *exploitative or predictive mode* in older adulthood (Brown et al., 2022; Spreng & Turner, 2021).

### Perspectives

Future work should aim to uncover the metabolic foundations of the SENECA model, in line with emerging frameworks positioning metabolism as the next frontier of aging research (Mattson, 2025; Pellerin, 2025; Shichkova et al., 2025), with efficient metabolic regulation constituting a central mechanism of an allostatic brain architecture (Theriault et al., 2025).

A promising direction lies in multimodal fMRI-PET approaches, which could yield a more comprehensive understanding of neurocognitive aging while mitigating vascular confounds (e.g., Palombit et al., 2022; Shokri-Kojori et al., 2019; Volpi et al., 2024; Wan et al., 2024). For example, functional MR spectroscopy could be used to quantify age-related changes in glutamate and GABA concentration, key neurotransmitters underpinning the excitatory-inhibitory (E/I) balance (Stanley & Raz, 2018). Notably, glucose hypometabolism has emerged as a critical factor in age-related declines in cognitive flexibility (Deery et al., 2024; Juttukonda et al., 2021; Tarumi & Zhang, 2018), with evidence of a similar inflection point in midlife (Huo et al., 2020; Thambisetty et al., 2013). Such metabolic shifts may partly reflect E/I imbalance, potentially linked to difficulties engaging GABAergic inhibitory processes linked to the DMN (Bi et al., 2020; Fortel et al., 2023; Xie et al., 2025), with disproportionate effects on functional brain activity in aging (Weistuch et al., 2021).

From an allostatic-first perspective, neuropathology could be reconceptualized as the outcome of allostatic regulation. Pathological cognitive decline may arise when cognition bears an excessive adaptive cost – a higher allostatic load than usual – in maintaining wiring economy under limited metabolic resources (i.e., hypometabolism). This hypothesis aligns with evidence that hypometabolism can precede clinical symptoms (Blázquez et al., 2022), highlighting the preventive potential of conceptualizing aging through the lens of energy regulation.

### Limitations

Caution must be taken regarding the cross-sectional design of the CamCAN database. Although estimations likely converge given a large enough sample (Sørensen et al., 2021), longitudinal data are required to confirm that the midlife represents a true transition independently from cohort effects (Cox, 2024). Notably, as observed in section 3.5, the cerebral substrates supporting the embodied semantic strategy varied substantially across participants. Age-invariant lifestyle factors, such as education, socioeconomic status, and physical activity, may help explain this variability, consistent with reports linking these factors to cognitive flexibility and language performance (Chung et al., 2012; Iso-Markku et al., 2024; Khalili et al., 2024; Perez et al., 2025; Rahman et al., 2023).

## Conclusion

How does the older adult brain reorganize to sustain cognitive flexibility?

By analyzing resting-state fMRI data from the population-based CamCAN database (N = 628; age 18-88), using a graph-theoretical framework grounded in structural balance theory, we found that midlife represents a critical transition period for brain network energy. During this phase, energy is reallocated along the sensory**-**transmodal hierarchy, shifting from higher-level networks (DMN–FPN) toward lower-level systems (SMN, CON, Auditory, Visual, Language). Crucially, despite this reallocation, global wiring economy remains preserved across the lifespan, suggesting an underlying allostatic mechanism or stability through change, likely regulated by anticorrelated dynamics. At the cognitive level, this supports a shift toward embodied semantic processing, wherein older adults rely on predictive, sensory-grounded strategies to sustain cognitive flexibility with reduced executive demands. In sum, our study provides a timely reconceptualization of healthy neurocognitive aging, framing it as an allostatic process, paving the way for extending the SENECA model to neuropathology.

## Data and code availability

All brain illustrations were made with the ggseg R-package (Mowinckel & Vidal-Piñeiro, 2020). Code for brain network energy computations and statistical analysis is made publicly available at: https://github.com/LPNC-LANG/SENECA_network_energy

## Supplementary information

Supplementary materials & results

## CRediT author statement

Conceptualization: CG; Methodology – Formal Analysis & Data Curation: CG; Methodology – Robustness Analysis; CG & SA, Writing – Original Draft: CG, MB; Writing – Review & Editing: CG, MB, MM; Funding Acquisition: MB, MM, SA; Supervision: MB, MM.

## Acknowledgments

This work was supported by the ANR project ANR-15-IDEX-02 and MIAI @Grenoble Alpes (ANR-19-P3IA-0003). This project has received financial support from the CNRS through the MITI interdisciplinary programs. The Cambridge Centre for Ageing and Neuroscience (Cam-CAN) research was supported by the Biotechnology and Biological Sciences Research Council (grant number BB/H008217/1).

