## Supplementary information for "Cognitive Flexibility and Brain Network Energy in Healthy Aging: an Allostatic Perspective from the SENECA model"

**Supplementary materials**

**Description of the 8 neuropsychological tests used to assess lexical production.** Please refer to original CamCAN articles for more information on behavioral datasets (Cam-CAN et al., 2014; Taylor et al., 2017).

***Cattell:*** *Cattell Culture Fair Test: Complete nonverbal puzzles involving series completion, classification, matrices, and conditions* (Cattell & Cattell, 1960)*.*

***Hotel Task:*** *Perform simulated tasks of a hotel manager: write customer bills, sort money, proofread adverts, sort playing cards, alphabetize a list of names. Total time must be allocated equally between tasks; there is not enough time to complete any task* (Shallice & Burgess, 1991)*.* ***In this study, we applied a log transformation and subtracted it from 1 to stay consistent with a decrease as age increases (i.e., 1-log(x)).***

***Picture Naming****: Name the pictured object presented alone (baseline), then when preceded by a prime object that is phonologically related (one or two initial phonemes), semantically related (low or high relatedness), or unrelated* (Clarke et al., 2013)*.*

***Proverb:*** *Read and interpret three English proverbs* (Huppert et al., 1994)*.*

***Sentence Comprehension:*** *Listen to and judge the grammatical acceptability of partial auditory sentences that begin with an ambiguous sentence stem (e.g., “Tom noticed that landing planes…”) followed by a disambiguating continuation word (e.g., “are”) in a different voice. Ambiguity is either semantic or syntactic, with empirically determined dominant and subordinate interpretations* (Rodd et al., 2010)*.*

***Story Recall:*** *Listen to a short story, recall freely immediately after, then again after a delay, and finally answer recognition memory questions* (Tulsky et al., 2003)*. Delayed recall measure used here.*

***Tip-of-the-Tongue (ToT):*** *Participants are asked to name famous faces and indicate if they know/don’t know/or have a ToT* (Brown & McNeill, 1966)*.* ***In this study, we subtracted the score from 1 to stay consistent with a decrease as age increases (i.e., 1-x).***

***Verbal Fluency:*** *Mean of letter (phonemic) fluency and animal (semantic) fluency task. For the phonemic fluency task, participants have 1 minute to generate as many words as possible beginning with the letter ‘p’. For the semantic fluency task, participants have 1 minute to generate as many words as possible in the category “animals”* (Lezak et al., 2012)*.*

**Supplementary results**

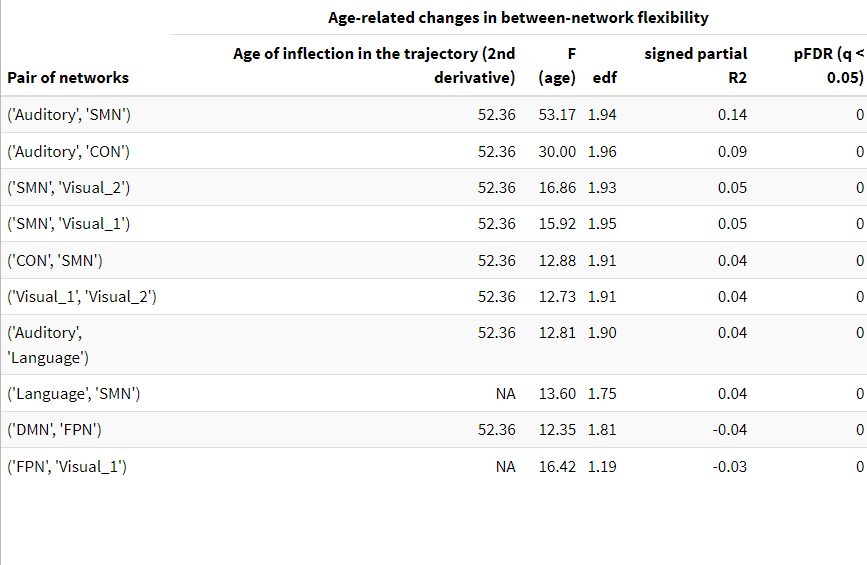

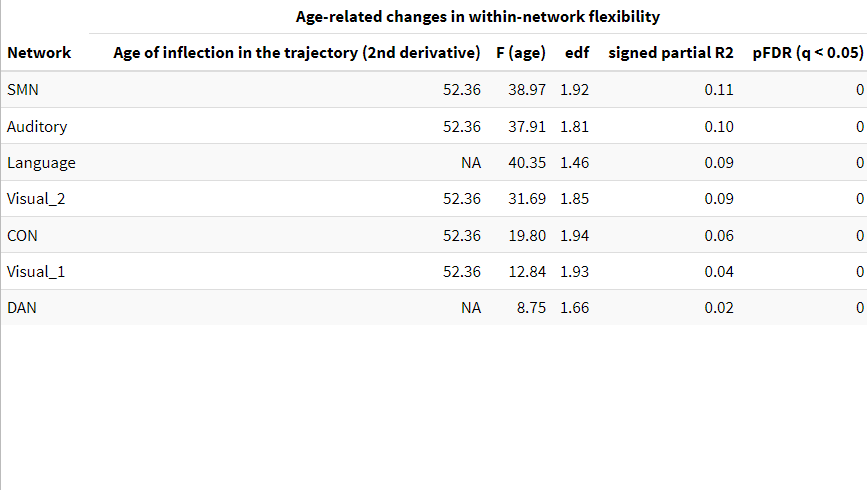

**Table S1. GAM statistics at the network level.** We report the significant trajectories with the 10 largest positive age-related effect

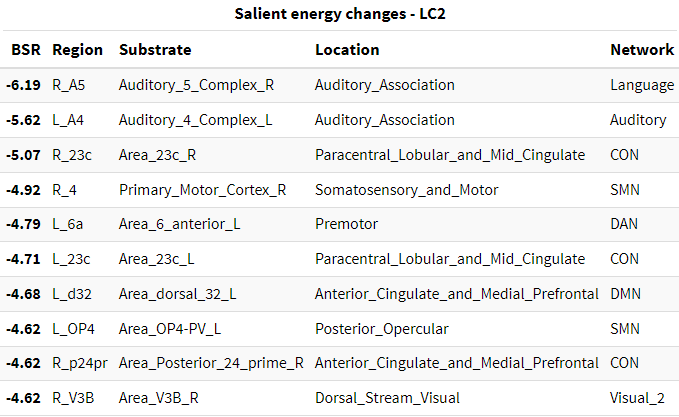

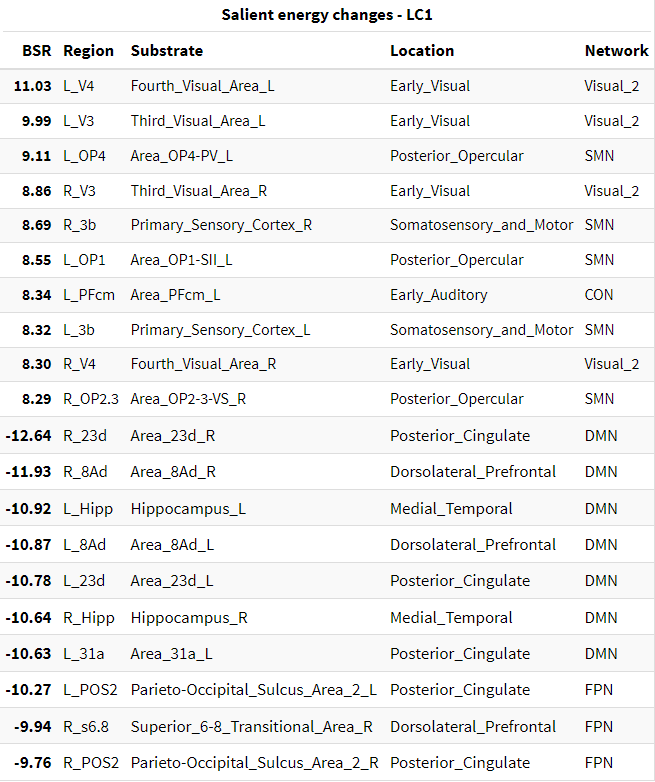

**Table S2. Salient features for each latent component of the PLS model.** We report the salient trajectories with the 10 largest negative and the 10 largest positive age-related effect

**Robustness analysis**

***Method 1:*** Following thresholding, null models preserving spatial and temporal autocorrelation (Shinn et al., 2023) preserves 70% (SD: 14%) of pairwise functional connections on average across all subjects. Significantly more connections were preserved in older adults, particularly beyond midlife (*F* = 67.29, *p* < .001, *edf* = 1.89, signed partial R² = - .17).

***Method 2:*** Following thresholding, the correlation screening approach (Lbath et al., 2024) preserves 18.4% (SD: 6%) of pairwise functional connections on average across all subjects, with no evidence for age-related variability (*p* = .55).

- **Whole-brain level**

We replicate the results reported in section 3.1.

- No change in flexibility across the lifespan with method 1 and 2 (ROPE fully covers the 89% HDI of posterior samples)
- Substantial changes in between vs. within network flexibility with method 1 (*F* = 25.33, *p* < .001, *edf* = 1.86, signed partial R² = - .07) and with method 2 (*F* = 73.91, *p* < .001, *edf* = 1.93, signed partial R² = - .19), with both exhibiting a similar inflection point at age 52.
- **Subnetwork level**

We replicate the results reported in section 3.2, observing similar changes in between-network and within-network flexibility across the lifespan.

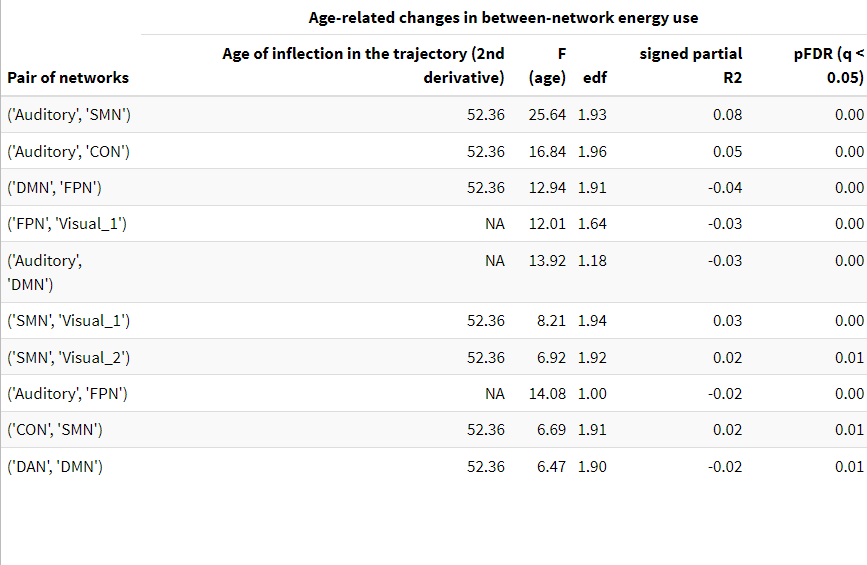

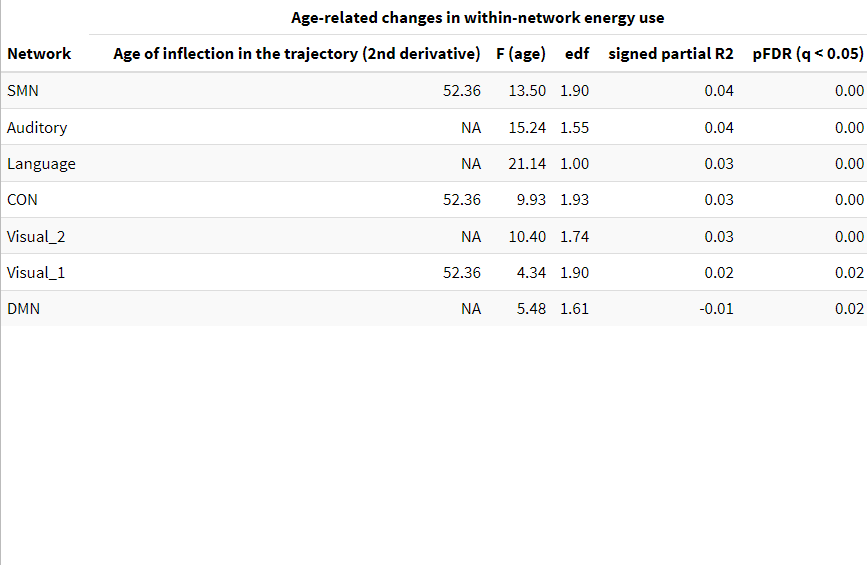

**Table S3. Thresholding method 1 - GAM statistics at the network level**

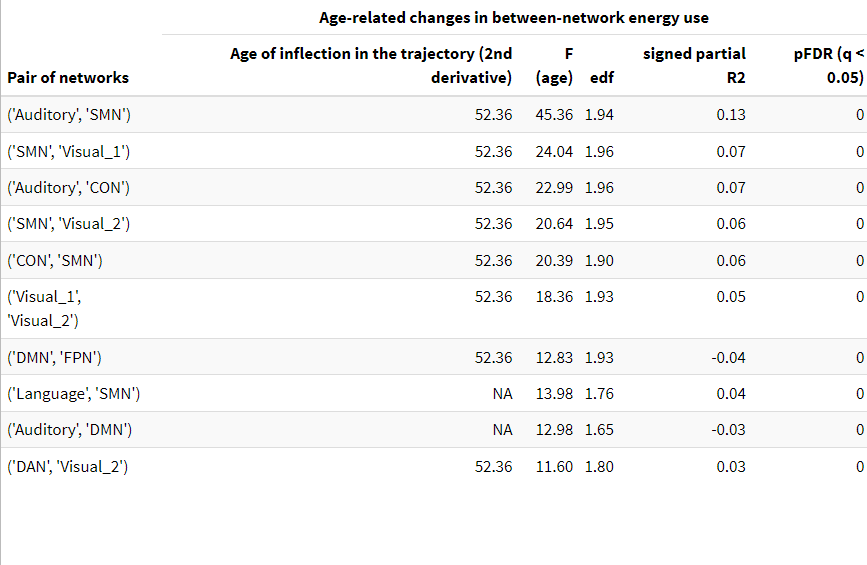

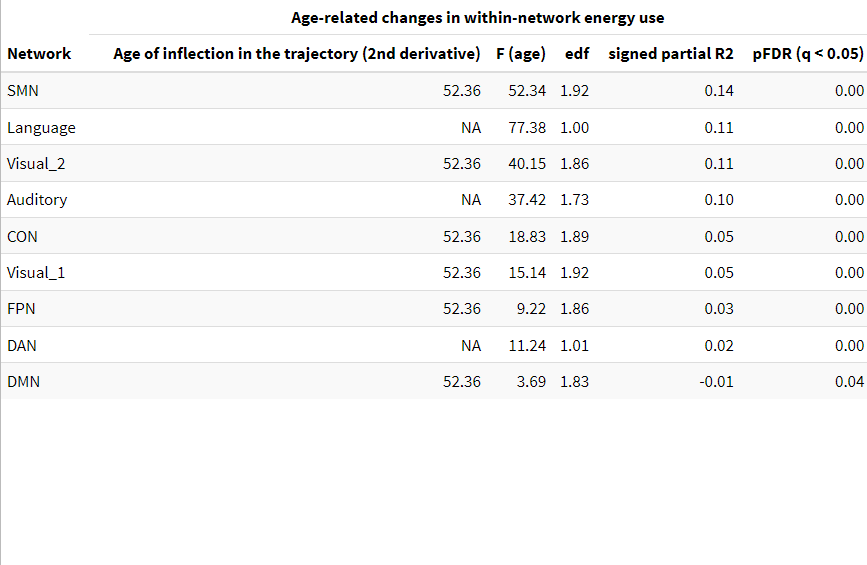

**Table S4. Thresholding method 2 - GAM statistics at the network level**

- **Region level**

As mentioned in the main text (section 3.4), region level analyses could not be reliably conducted after thresholding matrices using the correlation screening approach (Method 2). Only results for the null models (Method 1) are reported below.

We replicate the results reported in section 3.3, highlighting the PCC as a crucial DMN-FPN interface for flexibility, and its heterogeneous role within the DMN across the lifespan

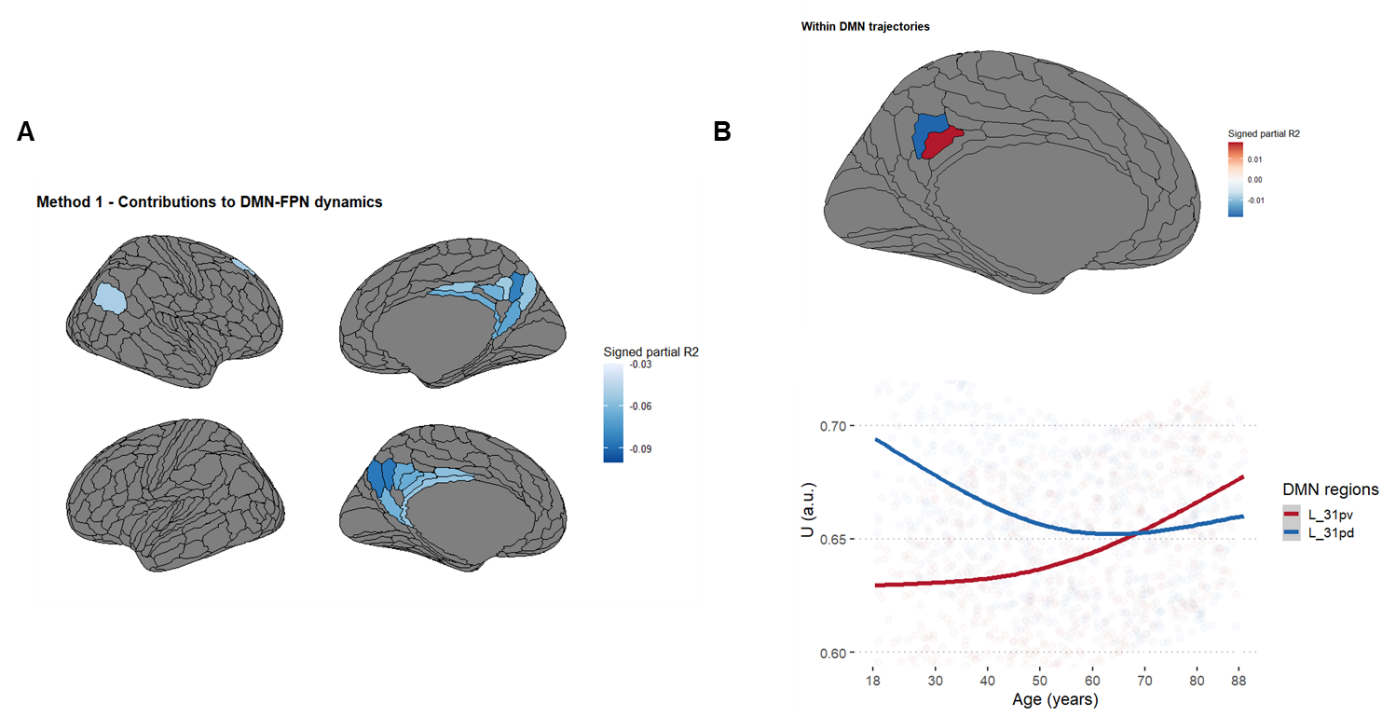

**Figure S1. Thresholding method 1 - Regions contributing to changes in (A) DMN-FPN and (B) within DMN flexibility**
